# Two-Photon Targeted, Quad Whole-Cell Patch-Clamping Robot

**DOI:** 10.1101/2022.11.14.516499

**Authors:** Gema I Vera Gonzalez, Phatsimo O Kgwarae, Simon R Schultz

## Abstract

We present an automated quad-channel patch-clamp technology platform for *ex vivo* brain slice electrophysiology, capable of both blind and two-photon targeted robotically automated patching. The robot scales up the patch-clamp singlecell recording technique to four simultaneous channels, with seal success rates for two-photon targeted and blind modes of 54% and 68% respectively. In 50% of targeted trials (where specific cells were required), at least 2 simultaneous recordings were obtained. For blind mode, most trials yielded dual or triple recordings. This robot, a milestone on the path to a true *in vivo* robotic multi-patching technology platform, will allow numerous studies into the function and connectivity patterns of both primary and secondary cell types.

## I. Introduction

Whole-cell patch-clamp is the gold-standard technique for obtaining high-fidelity electrical recordings of individual neurons. It has enabled the analysis of ion channel biophysics, membrane properties, cell excitability, post and presynaptic responses, neuronal interconnectivity and high order behavioural states, among many others. Despite providing high-quality data, the patch-clamp technique remains limited by inherently low throughput, steep learning curve and labour intensity.

The combination of patch-clamp automation and twophoton imaging enables the selection of specific cell types, thus presenting the opportunity to resolve single-cell characteristics with brain function by integrating anatomical, pharmacological and physiological data [1], [2]. As a result, hypotheses about the function of circuits involving specific cells or cell types during healthy and pathological states can be tested. This method, known as TPTP (Two-Photon Targeted Patching) [3], [4], directs pipettes filled with a fluorescent dye to patch cells that have been fluorescently labelled with an emission spectrum that allows them to be differentiated. Fluorescence can be induced via intracranial viral injection [4], breeding transgenic animals with cell-specific expression of fluorescent proteins [5], or intravenous injection of blood-brain-barrier-crossing viral vectors [6].

When attempting simultaneous recordings from multiple cells, the benefits of a robotic patch-clamp system may be even more significant [7]–[9]. Pairing patch-clamp recordings allows for precise measures of the connectivity between two neurons as it provides information on linked electrical activity at high temporal resolution, including subthreshold correlations which are not always possible to detect. Other methods, such as genetically encoded calcium indicators (GECIs) like GCaMP or voltage sensors, have not yet consistently shown comparable sensitivity permitting the identification of synaptic connections [12], [13]. In single-cell transcriptomics, paired recordings could also offer much more intricate biological detail [10], [11]. Transcriptomic changes underlying neuronal computation and development, for example, can be effectively investigated. Considering the challenges of carrying out multiple *in vivo* patch-clamp recordings [14]–[16], robotic automation appears to be the main solution for its widespread dissemination.

The use of the patch-clamp technique in live animal preparations has traditionally been limited to ‘blind’ recordings [17], [18]. In this technique, cells are detected by a change in impedance and the first cells encountered on the electrode path are selected as targets [9], [19]. Blind multi-patch-clamp has previously been automated *in vivo* [9]. Although these systems can greatly improve throughput and may be the only option for deep structures in the brain (*in vivo*), they have several disadvantages. Firstly, blind patching leads to a very reduced throughput for specific types of cells, such as interneurons, which are key players in brain function, but only account for approximately 15%-30% of the cortical cell population in rodents [20]. Therefore, in this kind of blind, automated multi-patching which skews towards statistically prevalent cells, the intra and interpopulation connections of secondary cell types cannot be studied. Not only that, but multi-patching without imaging can lead to a perceived false decrease in connectivity as this parameter is highly correlated to inter-somatic distance [16]; patching cells too far away will likely decrease the chances of observing a connection, while patching too close without image guidance can lead to the disruption of axons and dendrites of previously sealed cells with the next pipette, which could destroy the connections. Thus, detailed connectivity studies are not feasible with these systems. Moreover, since blind systems detect cells based on changes in impedance, they tend to patch the part of the cell membrane first encountered, rather than the centre, which has been linked to a decrease in yield [21]. Many issues can thus be solved by the use of two-photon microscopy to target fluorescently labelled neurons. For this reason, we have developed a robot that can do both blind and twophoton targeted multi-patch-clamping, extending the system of Annecchino et al. [1], which performed TPTP of single cells *in vivo*. The system has been tested in brain slices with success and will be adapted to *in vivo* multi-patching in the near future. We here present the results for both the blind and TPTP *ex vivo* modes.

## II. Methods

### A. Technological

The platform comprises a Ti:Sapphire laser (Newport/Spectraphysics MaiTai HP), a multi-photon microscope (Scientifica Ltd), and a 3 degrees of freedom (DOF) micromanipulator per channel (Sensapex or Scientifica) fitted with mechanical stability clamps (to improve pipette resistance against vibration, tissue deformation and pressure changes; similar to the ones in [25]). It incorporates patchclamp amplifiers, data acquisition (DAQ) hardware and four custom-developed pressure regulators, which are a refined and scaled up version from [1] and have a continuous pressure output range, as compared to discrete pressure systems in previous autopatchers (e.g. [9]). Control is via a custom-developed LabView program, which acquires frames directly from the microscope, thanks to the partial incorporation of SciScan into the software. The graphical user interface has been designed to help the user keep track of the different functionalities by having them in different modules: two-photon imaging, pressure control, manipulator control, electrophysiology control, etc. To help keep track of the four different channels/pipettes, they are colour-coded (Fig. 1).

**Fig. 1.**
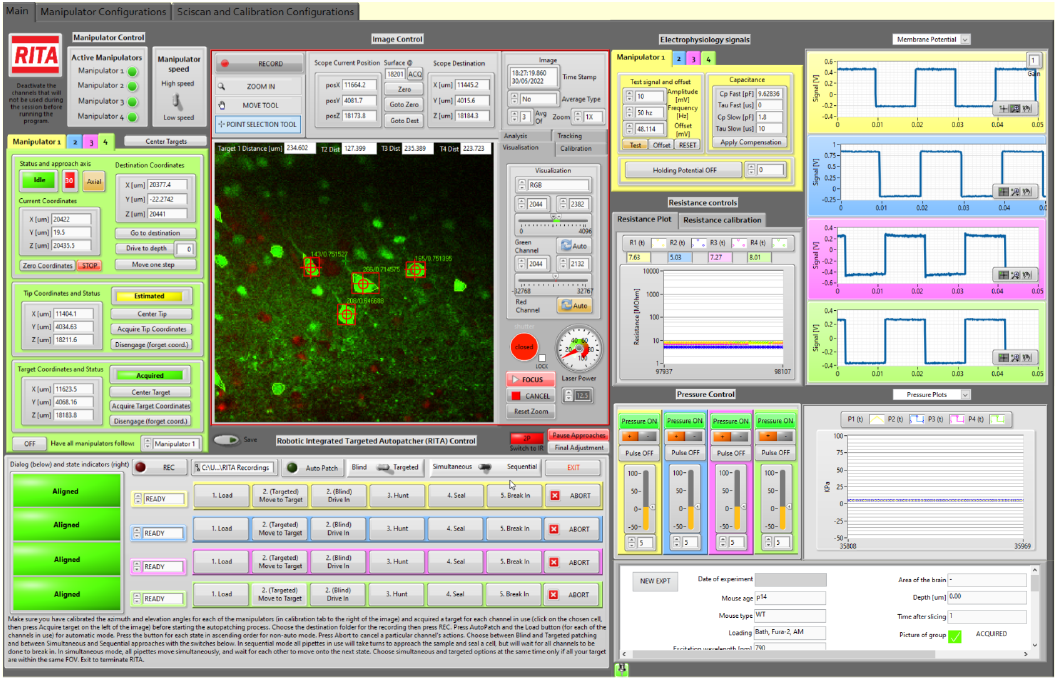
LabView control software user interface. Functionalities are separated into well defined modules and colour-coded according to channel for ease of use.

In targeted quad *ex vivo* mode (brain slices), a 3D stack of the target cells is acquired, and an image processing algorithm detects their centre of mass. The pipettes then simultaneously align in the x/y plane to a position where they have a direct path along the approach angle to their targets. This is followed by a single step movement along said axis, which leaves them in contact. At this point, the sealing protocol is activated. Upon successful seal, the user can choose to break-in or not, whereas break-in will automatically happen if set to auto mode. A twophoton (2P) image of a successfully patched group of four neurons is shown in Fig. 2.

**Fig. 2.**
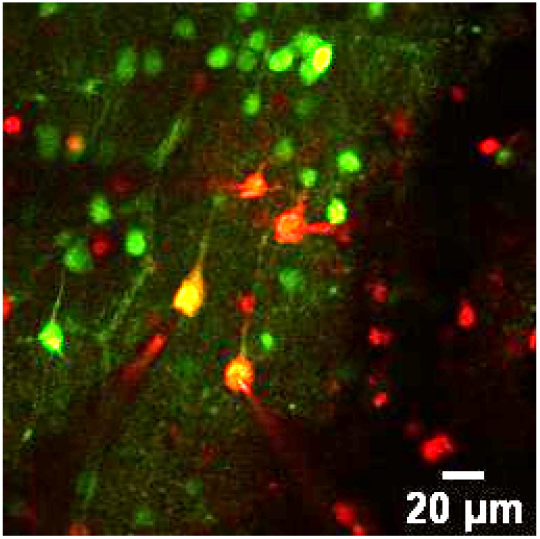
A two-photon image of four patched neurons. Cells were bath loaded with FURA 2-AM. Alexa 594 Hydrazyde dye (red) from the pipette internal solution diffuses into the cells as direct contact with their intracellular spaces is established, making the green cells yellow.

For blind quad *ex vivo* mode, the impedance of the pipettes is closely monitored to detect the presence of a cell near the tip as they move through the brain. When a cell is detected, movement in all other pipettes stops, and the sealing protocol is activated for the tip in question. Once a seal is achieved, the other pipettes resume the hunt, and when all have achieved a seal, break-in is activated in all the channels.

In TPTP mode, the two-photon images are processed by the software, segmented, and the centre of mass, area, contour and contrast of all present cells extracted in real time owing to their fluorescent print, using the same image-processing algorithm as [1]. It is the centre of mass that is used as the x/y target coordinates for the pipettes, so as to patch the centre of the cell, while the z-coordinate comes from the plane at which the cell is most in focus. This is calculated by maximizing the Contrast Focus Score (CFS) a parameter that is defined as the difference between the fluorescence of the background surrounding the cell and the fluorescence of the edges of the cell.

After each trial, pipettes are automatically cleaned in an enzymatic solution and then returned to their initial positions, allowing for reuse, following [22]. Clicking one button allows all channels to be offset, their voltage step functions to be activated and the pressure set to the required value, leaving them ready for the next trial. A ‘follow’ function is available to allow the user to move a single pipette and have the rest follow, both in the z and x/y coordinates, to move to a different brain location if needed. Alternatively, pipettes can be automatically sent to a certain depth. Temperature control for the perfusion system is also possible within the software through a Peltier system (Scientifica Ltd). The experimental notes section automatically records the output of each channel in each trial, input resistance, series resistance and membrane potential of successful seals and break-ins, as well as an image of the target cells, labelled with the channel that was used for each cell. This is complemented by the recording module, which automatically documents all other valuable information such as electrophysiology and pressure signals versus time, target coordinates, pipette coordinates and state of the pipette (hunting, sealing, etc) along time. Basic injection and connectivity protocols (Fig. 7) are also available to run within the program, making a complete software package for the basic needs of any experimenter, regardless of skill set.

### B. Biological

For targeted experiments, C57BL/6J mice of both sexes of postnatal ages 11-16 were bath loaded with FURA 2-AM according to the protocol by [23]: 2*µ*L of Pluronic F-127 (20% Solution in DMSO) and 13*µ*L of DMSO were added to the 50*µ*g of FURA 2-AM vial and the solution was vortexed and mixed with 2mL of resting artificial Cerebrospinal Fluid (aCSF). This solution was added to the loading chamber, for a slice incubation of 45 minutes at 35ºC. Slicing and resting aCSFs were prepared according to the same protocol. A laser excitation wavelength of 790nm was used in the 2P microscope for optimal visualization.

Imaging (for both targeted and blind) and slicing of blind experiments were done in aCSF with the following mM concentrations: 2 CaCl_2_ (Calcium chloride), 2.5 KCl (Potassium Chloride), 26 NaHCO_3_ (Sodium bicarbonate), 1.25 NaH_2_PO_4_ (Sodium phosphate monobasic monohydrate), 1 MgCl_2_ (Magnesium Chloride), 25 Dextrose and 125 NaCl (Sodium Chloride). Osmolarity was adjusted with dextrose to 338mOsm.

Internal solution for the pipettes was prepared in advance and frozen in aliquots with the following mM concentrations: 5 KCl (Potassium Chloride), 115 K-Gluconate (Potassium Gluconate), 10 HEPES (4-(2-hydroxyethyl)-1piperazineethanesulfonic acid), 4 Mg-ATP (Adenosine 5’triphosphate magnesium salt hydrate), 0.3 Na-GTP (Guanosine 5-triphosphate sodium salt hydrate) and 10 NaPhosphocreatine (Phosphocreatine disodium salt hydrate) respectively. pH was adjusted with KOH to reach 7.2-7.4 and sucrose was added to adjust the osmolarity to 310mOsm. 10mM Alexa 594 Hydrazyde was added on the day for pipette visualization.

## III. Results

### A. Quadruple targeted ex vivo patching

As a first step towards an *in vivo* two-photon targeted multi-patching robot, we developed an *ex vivo* mode making use of mouse brain slices. We achieved seal success rates of 54.2% (n=48) for targeted (Fig. 3) and 68.3% (n=82) for blind modes respectively (Fig. 5). This is in line with the success rates for manual patching but with far superior speed and throughput. An average of under 4 minutes was needed for all four channels to attempt seal and break-in in targeted mode, with total resistance averaging at around 65.4 *±* 4.75 MΩ and series resistance at 24.89 *±* 2.41 MΩ. In 50% of trials (Fig. 3), we obtained two or more simultaneous patches which could potentially be used for connectivity tests.

**Fig. 3.**
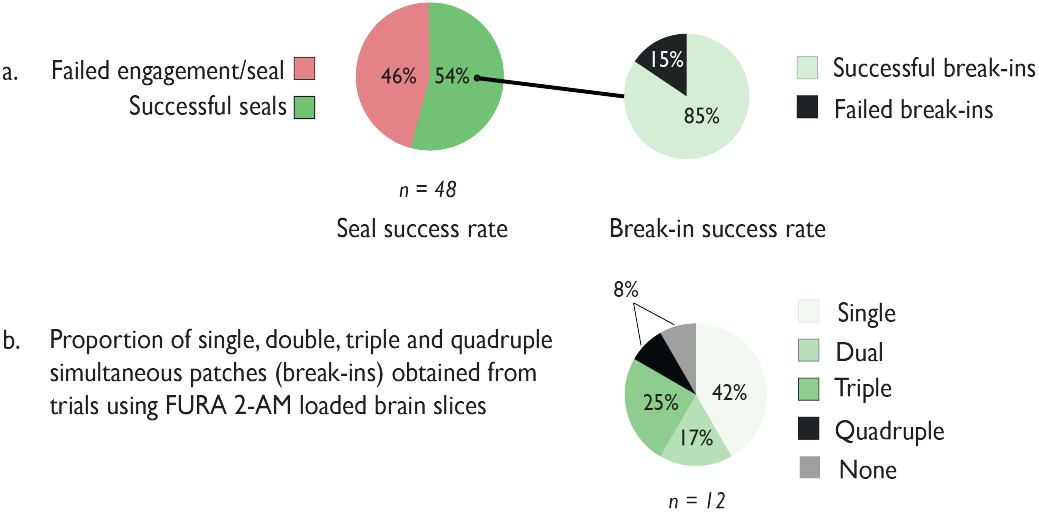
Success rates for *ex vivo* trials using dye-loaded (FURA 2-AM) brain slices. (a) Proportion of successful seal and break-ins.(b) The different number of simultaneous recordings obtained using four channels.

Example traces of the resistance and pressure signals (along time) of a successful quadruple patch are shown in Fig. 4.

**Fig. 4.**
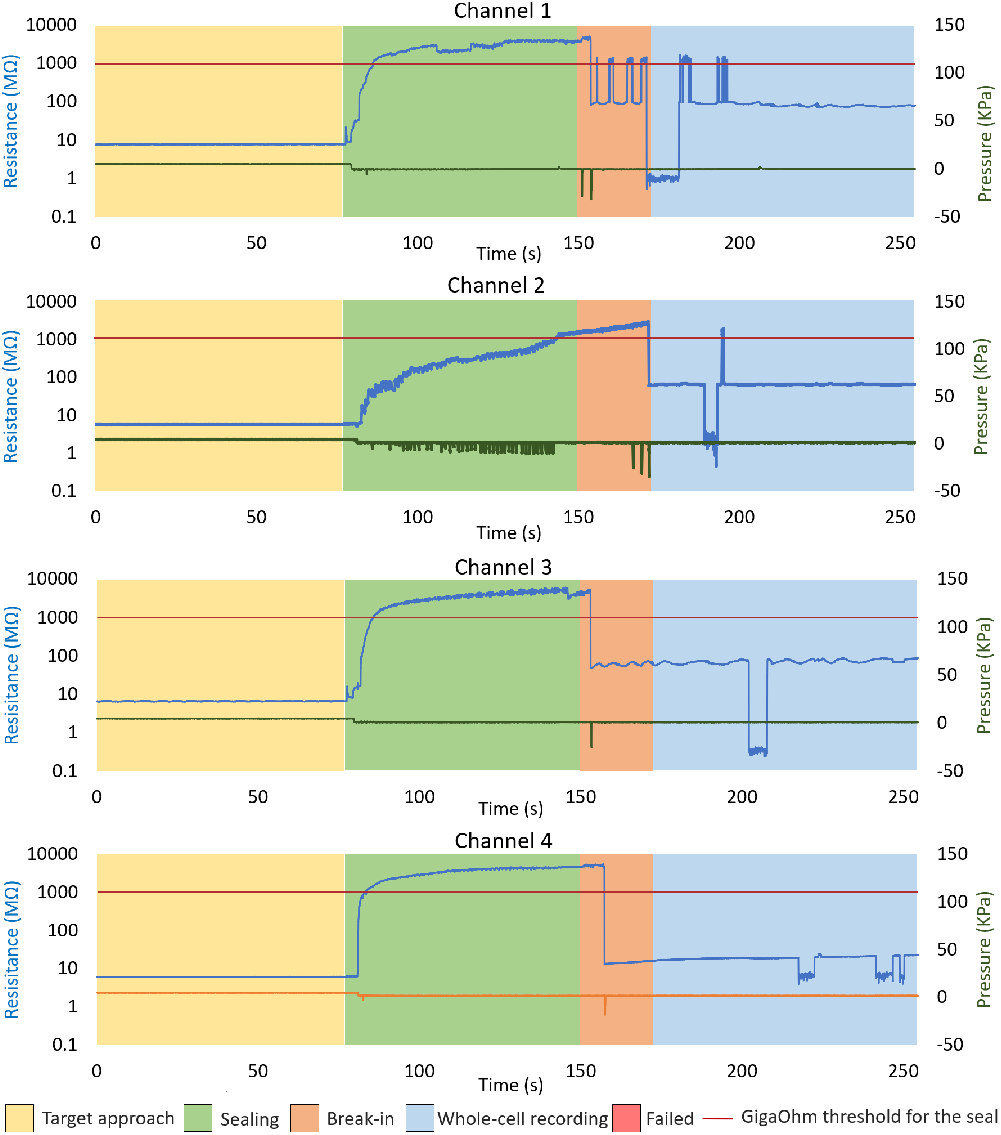
Stages of automated patching showing pipette resistance and pressure changes. Once the target cells are engaged, sealing automatically begins in all channels, releasing positive pressure and applying increasing suction pulses until a gigaseal is formed. Similar but stronger suction pulses are applied for break-in, which is also synchronized within all active channels. The square artifacts seen in the whole-cell mode arise from RITA changing to current clamp to check the resting membrane potential of the cell.

Our success rates are unfortunately not directly comparable to manual *ex vivo* patching or previous systems, as it is the first of its kind (multi-patch-clamping, two-photon guided, automated system). Upright bright-field microscopes with infrared (IR) light sources are the traditional method used in manual patch-clamp because it allows users to see cells dimpling, which is key to obtaining successful seals. On the contrary, dimpling is not seen in two-photon microscopy. Therefore, it would be expected of IR guided systems and manual users to achieve a higher success rate than a two-photon guided one. It should be noted, however, that our system still performed on par with previous *ex vivo* automated systems, like the work of Kolb et al. [29] (43% WCR for subcortical cells and 51% for cortical cells) or Koos et al. [30] (63.6% WCR for rat, 37% for human), despite these systems being IR guided and using only one manipulator. Our results are also slightly superior to the results presented by Wu et al. [28] (43.2% without manual intervention. IR guided but capable of matching up cells to two-photon fingerprints). Moreover, these systems cannot be expanded to be used *in vivo*, as ours can, due to the limited nature of IR imaging. Our results are also within the success rate range of manual users on IR systems, whose success is highly dependent on experience, falling anywhere between 30 and 80% [29].

Besides this, we cannot make a direct comparison with any of the automated systems for multi-patching, as the work of both [26] (brain slices, IR guided, 12 manipulators) and [27] (brain slices, IR guided, up to 10 manipulators) include manual steps in the algorithm that understandably increase yield when performed by experienced users. The work of Kodandaramaiah et al. [9] is also impressive, achieving 30.7% for blind triple or dual recordings in anaesthetized mice (in a system of 4 manipulators), a yield understandably lower than *ex vivo*, as the experimental paradigm is simpler.

### B. Quadruple ‘blind’ ex vivo patching

Fig. 7 shows typical current injection profiles that the program ran to characterise four patched neurons.

For blind mode patching in brain slices, we achieved 68.3% (n=82 including 14 quad and 8 triple trials) seal success rate for blind mode (Fig. 5). The average total resistance was 75.31 ± 6.03 MΩ and average series resistance was 23.50 ± 1.53 MΩ. Daily success rate varied from as low as 53% (n=17, 5 triple trials and 1 dual trial) to as high as 82% (n=19, 4 quadruple trials and 1 triple trial), we believe depending largely upon slice quality.

**Fig. 5.**
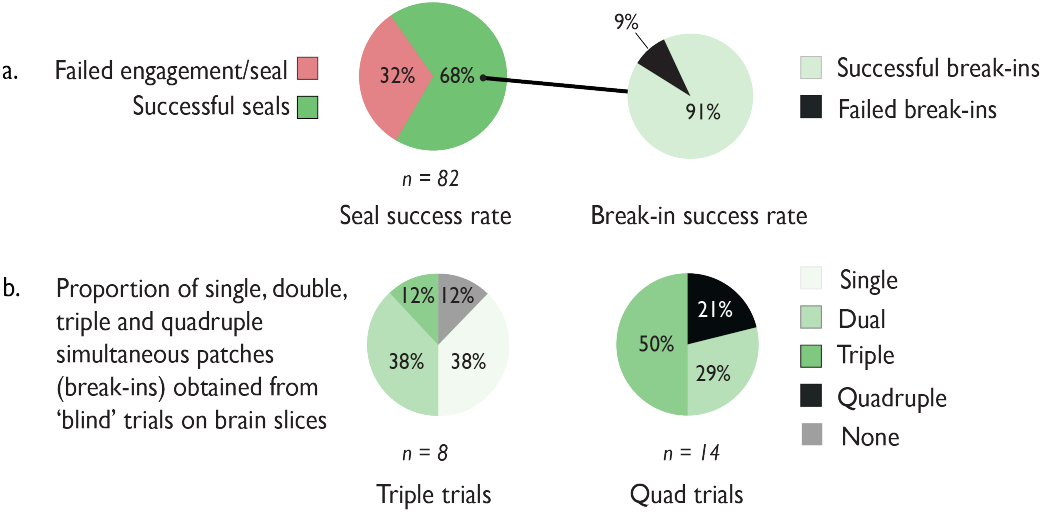
Blind patch success rate in brain slices. (a) Proportion of successful seal and break-ins. (b) The different number of simultaneous recordings obtained using three and four channels.

Example traces of the resistance and pressure signals (along time) of a triple patch are shown in Fig. 6, where it can be observed that Channel 4 failed to seal. This might point to either a false detection or to an unhealthy cell, as it was the first channel to detect (followed by Channels 3,2 and 1 in that order), and thus it was the most superficial cell. As can be seen in the figure, break-in is synchronized but sealing happens as soon as the cell is detected, and each channel maintains the seal until all active channels have attempted a seal.

**Fig. 6.**
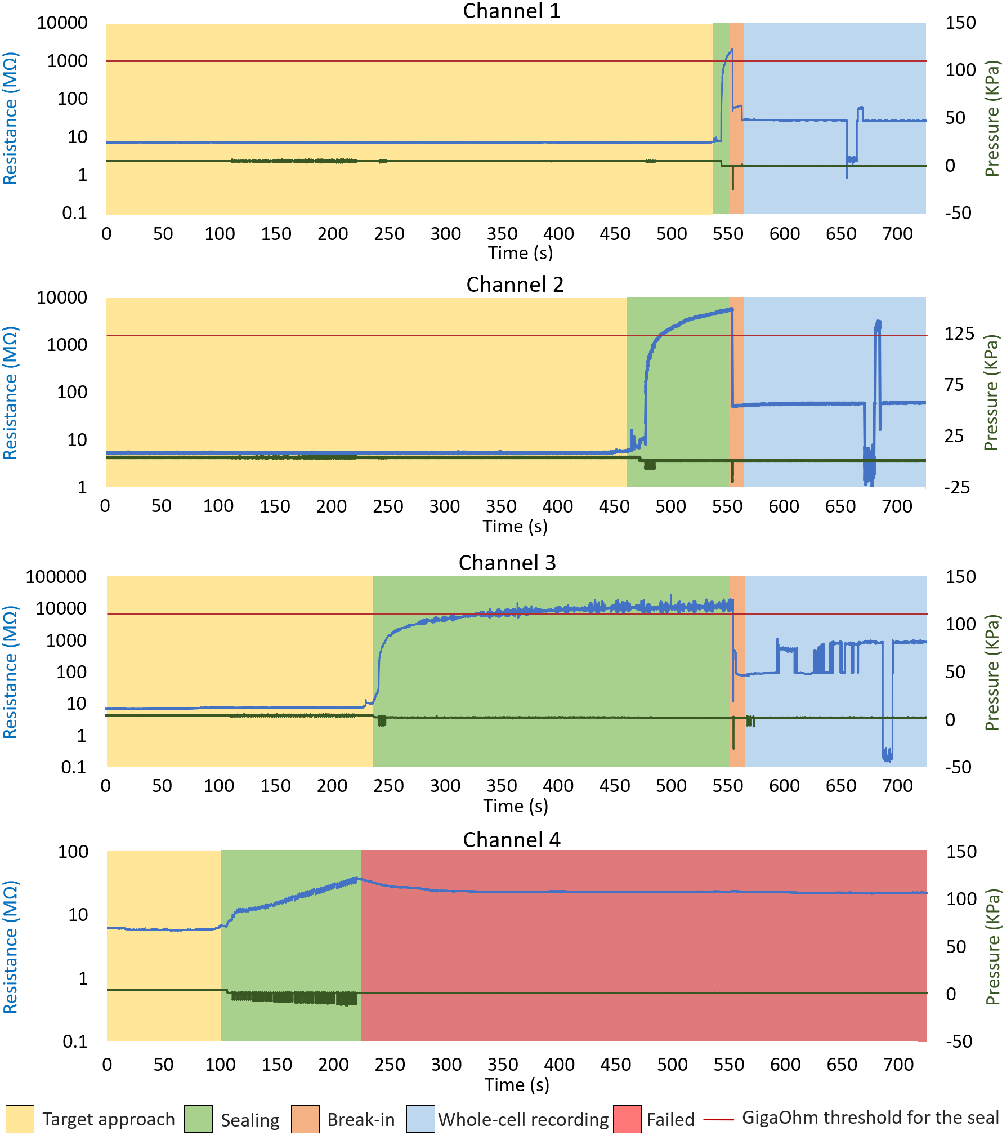
Stages of automated patching showing pipette resistance and pressure changes. Once the target cells are engaged, sealing automatically begins in all channels, releasing positive pressure and applying increasing suction pulses until a gigaseal is formed. Similar but stronger suction pulses are applied for break-in, which is also synchronized within all active channels.

**Fig. 7.**
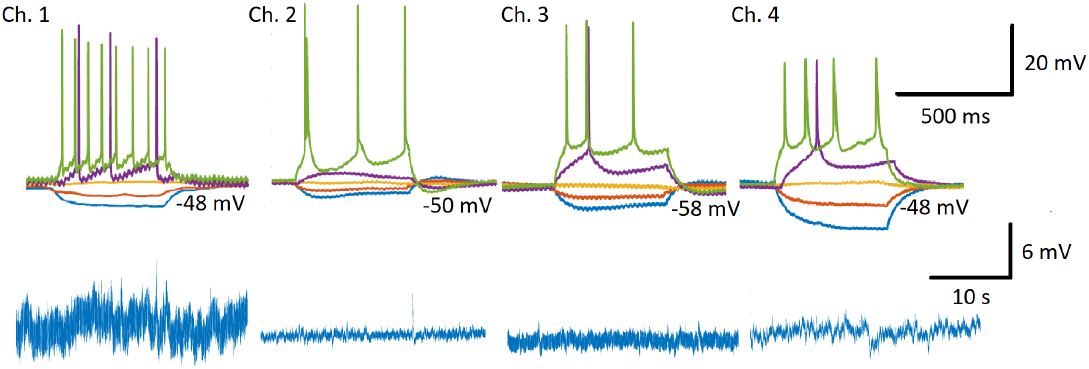
(Top) Example of the responses of four ‘blindly’ patched cells in response to the current pulses applied through the injection protocol module, following successful patching. 500 ms−long pulses from −100 to +280 pA in 20 pA steps were applied. Here, responses for -40, -20, 0, 20 and 60 pA pulses, are shown. (Bottom) Recordings of the same cells with no injections.

Based on these promising statistics, we envisage that our robotic platform will help make multiple patch-clamp electrophysiology accessible to a broader range of laboratories across the world. Additionally, it might be used as an aid by experienced patch-clamp researchers, either by using the pressure module to avoid mouth suction where there may be health safety concerns, or by saving time in numerous ways: automatically recording data, synchronised movement of pipettes, automated cleaning of pipettes for reuse, etc. It will thus make thousands of new neuroscience studies possible and more efficient.

## IV. Conclusion

Robotic automation helps mitigate low patch-clamp success rates and offers several benefits, including faster skill assimilation for new operators, standardized recording quality, and improved throughput. It also enables scaling up of the technique to simultaneous patch-clamp recording of multiple cells *ex vivo* and *in vivo*, enabling assays of synaptic coupling and many new research lines in basic and translational neuroscience. With further increase of degree of automation to include other aspects of experimental workflow, such as craniotomies [24], there is the potential for further reduction in human-derived experimental variability.

Our targeted quad patch-clamp system allows scalable and reproducible electrophysiology studies to be conducted across a variety of laboratory settings, offering for the first time, robotically automated recording of subthreshold signals from multiple genetically and optically targeted cells in tandem. As we have made use of only information available *in vivo*, we envisage straightforward extension of the platform from *ex vivo* to *in vivo* application.

## Acknowledgment

We thank J Sjöström and his laboratory for useful discussions on multiple patch-clamp recording, and L Annecchino for useful discussion related to his previous work on the platform.

